# Evaluation of intron-1 of odorant-binding protein-1 of *Anopheles stephensi* as a marker for the identification of biological forms or putative sibling species

**DOI:** 10.1101/2021.12.03.470951

**Authors:** Om P. Singh, Shobhna Mishra, Gunjan Sharma, Ankita Sindhania, Taranjeet Kaur, U. Sreehari, Manoj K. Das, Neera Kapoor, Bhavna Gupta

## Abstract

**Background:** *Anopheles stephensi*, an invasive malaria vector, has been reported to have three biological forms identifiable mainly based on the number of ridges present on the egg’s floats. Recently, the first intron of the odorant-binding protein-1 (*AsteObp1*) has been introduced as a molecular marker for the identification of these forms, and based on this marker, the presence of three putative sibling species (designated as species A, B and C) has been proposed. However, there is no data on the association of proposed markers with biological form or putative species on field populations.

**Methods:** Field collected and laboratory-reared *An. stephensi* were characterized for biological forms based on the number of ridges on the egg’ s float. DNA sequencing of the partial *AsteObp1* gene of *An. stephensi* individuals were performed by Sanger’s method, either directly or after cloning with a plasmid vector. Additionally, *AsteObp1* sequences of various laboratory lines of *An. stephensi* were retrieved from a public sequence database.

**Results:** *AsteObp1* intron-1 in Indian *An. stephensi* populations are highly polymorphic with the presence of more than 13 haplotypes exhibiting nucleotides as well as length-polymorphism (90-to-121 bp). No specific haplotype or a group of closely related haplotypes of intron-1 was found associated with any biological form identified morphologically. High heterozygosity for this marker with a low inbreeding coefficient in field and laboratory populations indicates that this marker is not suitable for the delimitation of putative sibling species, at least in Indian populations.

**Conclusions:** *AsteObp1* cannot serve as a marker for identifying biological forms of *An. stephensi* or putative sibling species in Indian populations.

## Background

*Anopheles stephensi* is an efficient malaria vector responsible for malaria, mainly in urban areas. Earlier, this species was reported to be distributed mainly in countries of the Middle East and South Asia [1]. During the last few decades, the distribution of this species has expanded to new geographical localities, such as Lakshadweep island of India [2], Sri Lanka [3], countries of the Horns of Africa [4-8] and the Republic of Sudan [6, 9], and have been reported to transmit malaria parasite at least in Djibouti, Horn of Africa [4, 7, 10]. A high probability of their presence within many urban cities across Africa has been predicted [11], which poses a major threat to the elimination of urban malaria from the African continent. In consequence of recent invasions of this species in several countries, the World Health Organization issued an alert to affected and surrounding countries to take immediate actions [6].

*An. stephensi* in India is predominantly found in urban areas due to the presence of favourable breeding habitats in urban settings and is considered an urban malaria vector. In the 1940’s, Sweet and Rao [12] and Rao et al. [13] described the presence of two races, i.e., ‘type form’ and ‘*var. mysorensis’* in the *An. stephensi* based on the dimension of eggs, length of eggs-floats and numbers of ridges on the float. The ‘type form’ was reported mainly from the urban area and ‘*var. mysorensis*’ from the rural area [12, 14]. Initial reciprocal crossing experiments showed a certain degree of sterility between these two races [15] but subsequently, it was demonstrated that there is no reproductive barrier between these two races (hereafter, described as ‘biological form’) and the proposed status of subspecies was refuted by Rutledge et al. [16]. Later, an additional form “intermediate” was designated by Subbarao et al. [14] based on genetic evidence derived from crossing experiments between the ‘type form’ and ‘*var. mysorensis*’.

The ‘type form’ is reported to be an efficient malaria parasite vector, while ‘*var. mysorensis*’ is considered to be poor in malaria transmission [14], though the latter is reported to be a vector in Iran [17-18]. Laboratory feeding experiments suggest differences in the success of sporogonic development of rodent malaria parasites in the biological forms [19-20]. Due to the suggested differential epidemiological implications of biological forms, it is desirable to determine their biological forms in all epidemiological studies. The classical method for the identification of biological forms of *An. stephensi* involves morphometrics of eggs, which is a cumbersome process; for which, live female mosquitoes are to be transported to the laboratory and their eggs have to be obtained for the counting of ridges on the egg’s floats. Nagpal et al. [21] reported that the ‘spiracular index’ (ratio of the length of anterior spiracle with the length of the thorax) of *An. stephensi* can be used for the discrimination of biological forms in the adult mosquitoes, which was higher in ‘type form’ as compared to ‘*var. mysorensis*’ in Rajasthan, India (an arid zone). They also reported that the two forms of *An. stephensi*, i.e., ‘type form’ and ‘*var. mysorensis*’, occupy different ecological niches where ‘*var. mysorensis*’ prefers outdoor habitat and ‘type form’ indoors [21]. The ‘spiracular index’, which measures the degree of spiracular opening, may serve as an adaptation to water-loss regulation, and therefore needs to be evaluated in non-arid zones also. Recently, a molecular marker was introduced by Gholizadeh et al. [22] for the identification of all three biological forms. This group sequenced the odorant-binding protein-1 (*AsteObp1*) gene of *An. stephensi* from four laboratory strains, two strains representing ‘*var. mysorensis’* and one of each representing ‘type form’ and ‘intermediate’. Based on sequences of a total of 20 samples from all of the four colonies, they demonstrated fixed differences (no heterozygotes were reported) in the first intron of the gene among three biological forms. This prompted them to propose the *AsteObp1* intron-1 as a molecular marker for the identification of biological forms. However, this marker was not tested on field populations. Subsequently, the same research group [23], speculated the three biological forms to be distinct sibling species and designated them as species A, B and C, which correspond to ‘type form’, ‘intermediate’ and ‘*var. mysorensis’*, respectively. However, they didn’t establish an association of the *AsteObp1* intron-1 haplotypes with the eggs’ morphology. In the absence of any study showing the association of *AsteObp1* haplotypes with biological form in field populations, we investigated the extent of polymorphism in *AsteObp1* intron-1 in Indian *An. stephensi* (field populations as well as laboratory colonies) and validated their association with biological forms or proposed sibling species.

## Material & methods

### Mosquito sampling and processing

*Anopheles stephensi* samples were collected from urban, peri-urban and rural localities. The list of localities and their geographical coordinates is provided in **Table 1**. The resting mosquitoes were collected from dwellings in the morning between 6:00 and 8:00 AM with the help of a mouth aspirator and flash-light. The live mosquitoes were carefully transferred in a thermocol box and transported to the insectary maintained at 27 °C and 70-75% RH. Mosquitoes were provided access to soaked raisins and water-pads. Mosquitoes were allowed to attain the gravid stage. Gravid mosquitoes were provisionally identified for species using a magnifying glass and individual *An. stephensi* were transferred inside separate paper cups containing a small amount of water and having the inner side lined with blotting paper. The mosquitoes were kept in the cup overnight to allow them to lay their eggs. The next morning, female mosquitoes which laid their eggs were removed, anesthetized with diethyl ether, and confirmatory identification of *An. stephensi* was performed under a binocular microscope following keys by Christophers [24]. The mosquitoes were preserved in individual microcentrifuge tubes containing a few drops of isopropanol for molecular studies. The eggs of corresponding mosquitoes laid on filter paper were removed onto a glass micro-slide and the number of ridges present on one side of eggs’ float was scored following Rao et al [13] from 10 to 15 eggs under a stereo-binocular microscope using a 10X objective lens. Finally, the mode number of ridges was determined for each egg batch. Mosquitoes from laboratory colonies of *An. stephensi* originating from New Delhi (NIMR strain) and being maintained since the year 2011, was also characterized for the number of ridges present on egg’s float. DNA was isolated from individual mosquitoes manually following methods by Black and Duteau [25]. Besides, genomic DNA isolated from *An. stephensi* individuals from a laboratory colony (F5) originating from Chennai (urban area) and *An. stephensi* strain STE2 (origin, Delhi, India, supplied by BEI Resources, NIAID, NIH, USA) were also used for molecular studies. A total of 143 samples were included in this study, out of which 137 samples were sequenced successfully and 119 samples were successfully characterized for the number of ridges on eggs, of which 27 were characterized as ‘*var. mysorensis’*, 30 as ‘intermediate’ and 62 as ‘type form’.

**Table 1.** Geographical locations of *Anopheles stephensi* collection sites and other laboratory trains used for molecular studies.

| Populations/strains | Geographical coordinates | Urban/rural | Bio-types |
| --- | --- | --- | --- |
| **Wild populations** |  |  |  |
| 1. Bengaluru Karnataka state | 13°00' N, 77°63' E | Urban | Type form |
| 2. Karamcnand Pur, Haryana | 28°04' N, 77°13' E | Rural | Intermediate and 'var. mysorensis' |
| 3. Jaisingh Pur, Haryana | 28°04' N 77°14' E | Rural | Type forms, 'intermediate' and *var. mysorensis* |
| 4. Chilawali, Haryana | 28°17' N, 76°95' E | Rural | *var. mysorensis* |
| 5. Ghummanhera South-West Delhi district | 28°53' N, 76°92' E | Peri-urban | Intermediate and *var. mysorensis* |
| **Laboratory colonies** |  |  |  |
| 1. NIMR strain (Delhi) | 28° 38 N', 77° 12' E | Urban | Type form |
| 2. Chennai | 13° 07 N', 80° 23' E | Urban | Type form |
| 3. STE2 strain (Delhi) | 28° 38 N', 77° 12' E | Urban | Not known |

### PCR amplification, cloning and sequencing

Initially, the amplification of partial *AsteObp1* was performed using primers OBP1F1 (5’ - CGT AGG TGG AAT ATA GGT GG-3’) and OBP1R2 (5’ -TCG GCG TAA CCA TAT TTG C-3’ [22] which covers full intron-1 and partial intron-2. PCR was performed with Hot Start Taq 2X Master Mix (New England Biolabs Inc) in a 20 µL reaction mixture containing 1.5 mM of MgCl_2_, 0.5 units of Taq polymerase and 0.25 µM of each primer. The PCR cycling conditions were: a denaturation step at 95 °C for 3 min, 35 cycles, each with a denaturation step at 95 °C for 30 s; annealing at 55 °C for 30 s and extension at 72 °C for 45 s, followed by a final extension at 72 °C for 5 min. PCR products were cleaned using ExoSAP-IT™ (ThermoFisher Scientific) and sequenced on both strands of DNA using ABI BigDye Terminator v3.2 (ThermoFisher Scientific) following the manufacturer’s protocol.

To phase out haplotypes in heterozygotes, the PCR products of 12 heterozygous individuals exhibiting different patterns of mixed bases in sequence chromatograms were selected for cloning followed by sequencing. For cloning, PCR products were amplified using a high-fidelity Taq DNA polymerase to avoid or minimize PCR errors. Briefly, PCR reactions were performed in a total volume of 25 μL containing 0.5 units of Phusion (New England Biolabs) Taq DNA polymerase, Phusion HF reaction buffer (containing 1.5 mM MgCl_2_ and 200 mM of each dNTP), 0.5 μM of each primer and 0.5 μL of PCR product. PCR thermal cycling conditions were: initial denaturation at 98 °C for 30 s, 35 cycles, each of denaturation at 98 °C for 10 s; annealing at 65 °C for 30 s and extension at 72 °C for 45 s, followed by a final extension at 72 °C for 2 min. Five μL of PCR product was loaded onto 2% agarose gel and visualized under UV illumination. The remaining PCR products were purified using the QIAquick PCR purification kit (Qiagen Inc) following the vendor’s protocol. The purified PCR product was ‘A’-tailed using dATP in the presence of Taq polymerase. The reaction mixture (50 μL) contained 0.25 μL of Taq polymerase (KAPA Biosystem, USA), 5μL of 10X buffer containing 1.5 mM MgCl_2_, 5μL of purified PCR product, 1μL of 10mM dATP and incubated at 70 °C for 20 min. Three μL of purified A-tailed PCR product was then ligated into pGemT®-T vector (Promega Corporation) in a 1:3 vector to insert ratio following the recommended protocol by the manufacturer. Then 5 μL of the ligated product was transformed into DH5-alpha competent E. coli (New England Biolab) and 90 μL of the transformed product was spread onto Ampicillin-IPTG/X-Gal LB agar plates and incubated overnight at 37 °C for blue/white colony screening. Plasmid DNA from five clones from each mosquito was PCR amplified and sequenced from both directions of the DNA strand as described above.

For subsequent sequencing reactions for the characterization of full intron-1 (short-PCR), we designed a different primer set (OBP_INTF: 5’ -CGC CGT GAT GCC GAA TA-3’ and OBP_INTR: 5’ -ATT GTC GTC CAC CAC CTT G-3’) from flanking exons. The PCR conditions were similar to what we used for the PCR amplification for direct sequencing using primers OBP1F1 and OBP1R2, except for the duration of extension time in PCR-cycling, which was reduced to 30 s. A total of 137 samples, which includes 109 samples characterized for egg ridges count were successfully sequenced for *AstObp1-*intron-1.

### Haplotype phasing in heterozygotes

Due to the presence of indels at multiple loci in the intron-1, the phasing of haplotypes in heterozygous sequences was performed using the online tool Poly Peak Parser available at http://yosttools.genetics.utah.edu/PolyPeakParser [26]. The haplotypes identified through sequencing of cloned samples, homozygotes and heterozygotes for single SNP were taken as reference haplotypes. While using software signal ratio cut-off value was adjusted below the level of the true peaks, but always above the noise level. Any new haplotype determined by the software which was not present in reference sequences was treated as ‘unidentified’ because of the uncertainty of phase.

### Genetic distance, haplotype networking and phylogenetic analyses

All the haplotypes of intron-1 of *An. stephensi* and *An. gambiae* (GenBank accession AY146721) were aligned using MUSCLE implemented in the application MEGA7 [27]. The sequence of *An. stephensi* was treated as an outgroup. The genetic distances among all haplotypes of *An. stephensi* and phylogenetic analyses were performed using the T92 model (Tamura 3-parameter), the best model as determined by MEGA7 based on the lowest Bayesian Information Criterion (BIC) scores. The bootstrap consensus tree inferred from 500 replicates was taken to represent evolutionary history. Haplotype network was constructed using NETWORK 10 (https://www.fluxus-engineering.com/sharenet.htm). The pairwise differences between and within biological forms, and *F*_ST_ between biological forms, based on the *AsObp1* haplotypes were estimated using software Arlequin [28],

### Retrieval of *AsteObp1* intron-1 sequences from publicly available genome database of laboratory strains of *An. stephensi*

Sequences of the intron-1 of *AsObp1* were retrieved from publicly available genome sequence databases of various strains of *An. stephensi*, i.e., AsteI2 strain (origin: India), SDA500 strain (origin: Pakistan) Hor strain (origin: India) and UCISS2018 strain (probably Indian origin), along with information on their biological forms, if available.

## Results

### Characterization of *AsteObp1* intron-1 haplotypes

A total of nine haplotypes (H1 to H9) were identified through sequencing of cloned PCR products from heterozygous individuals (**Figure 1**). Direct sequencing of the short-PCR product covering intron-1 revealed the presence of an additional six haplotypes (designated as H10 to H15) based on either homozygous sequences or heterozygous sequences with polymorphism at a single nucleotide base. Considering only intron-1 (the region proposed as a marker for the discrimination of biological forms or sibling species by Gholizadeh et al., 2015 and Firooziyan et al., 2018), there were 13 haplotypes because haplotype H1 was identical to H7, and H4 was identical to H5 in the intron-1 region. Thus, there were 13 haplotypes with respect to intron-1. Indels were present in both introns, but intron-1 exhibited a higher degree of length polymorphism which varied between 90 and 121 bp (90, 114, 116, 120 and 121 bp). The nucleotide sequences of all haplotypes are available at GenBank (accession nos. ON564278-ON564292)

**Figure 1.**
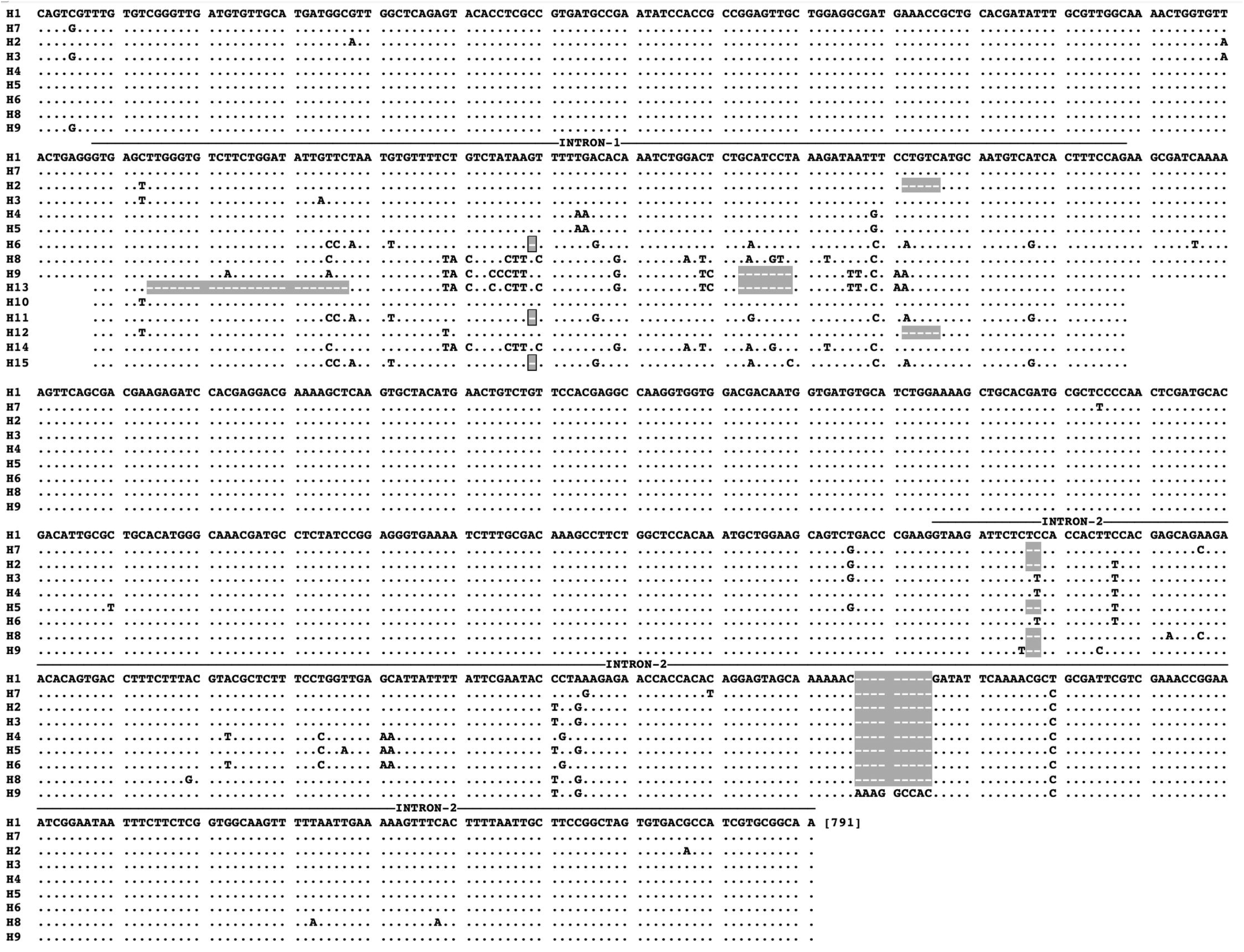
Sequence alignment of partial *AsteObp1* haplotypes covering intron-1, identified in this study. Dots (.) represent nucleotide sequence identical to first row of sequence and dashes (’ -’) represent gap due to indel (highlighted with gey colour).

### Distribution of haplotypes in different populations

The distribution of haplotypes in laboratory colonies and field populations is shown in **Table 2**. Haplotypes in nine heterozygous sequence reads could not be phased out due to the presence of additional haplotypes, which require additional cloning and sequencing for phasing. The *AsteObp1* intron-1 exhibited extensive polymorphism in field populations. A high degree of heterozygosity, as per expectation, was observed in field populations, i.e., in Karamcand Pur (*H*_O_=87%, *H*_E_=87%), Jaisingh Pur (*H*_O_=86%, *H*_E_=83%) and Bengaluru (*H*_O_=83%, *H*_E_=79%) (**Table 2**). The laboratory colony (NIMR strain) which has been maintained in the laboratory since 2011, had only two haplotypes (*H*_O_=46%; *H*_E_=45%). In all cases, the inbreeding coefficient was close to zero (−0.05 to 0.00), suggesting random breeding. In populations with a sample size of less than 10, we didn’t calculate heterozygosity and inbreeding coefficient, but the majority of the individuals in such populations were heterozygous (**Table 2**).

**Table 2.**
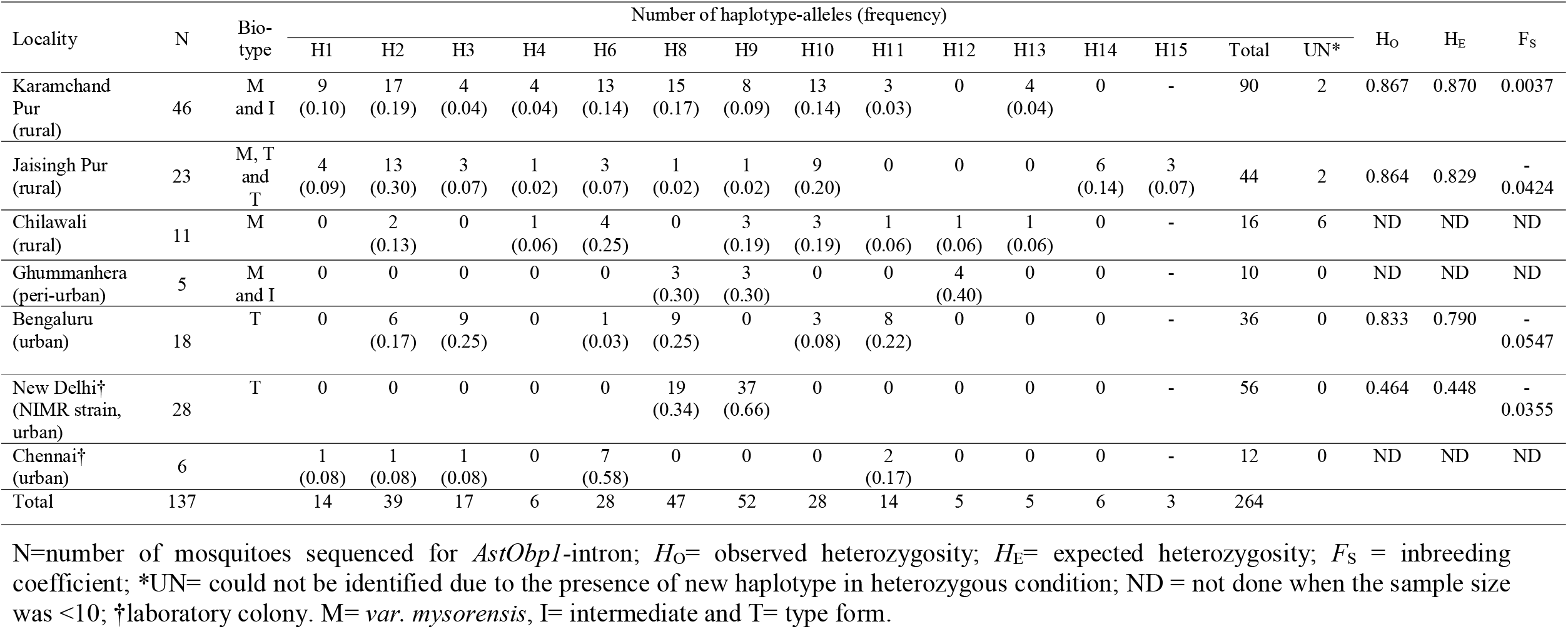
Distribution of *AsteObp1* intron-1 haplotypes in different population.

### Genetic distances, phylogenetic analyses and haplotype networking

To infer genetic relatedness among different haplotypes, haplotype clustering was performed based on (i) pair-wise genetic distance; (ii) phylogenetic tree, and (iii) minimum spanning haplotype network. For haplotype clustering, 13 haplotypes obtained in this study were analyzed together with five *AstObp1* intron-1 haplotypes identified by Gholizadeh et al. [22] and Firooziyan et al. [23] (Supplementary **Table S1**)

Pair-wise genetic distance between haplotypes, as estimated using Timura 3 Parameter (T3P) nucleotide substitution model, has been displayed in Supplementary **Figure S1**). Based on the genetic distance, haplotypes can be clustered into four groups: (i) the six haplotypes, H1, H10, H2, H3, H12 and H4 were genetically closer, with genetic distances ranging from 1-4%; (ii) H6, H11 and H15 were closer, with genetic distance ranging between 1-2%; (iii) H8 and H14 were closely related with a distance of 1%, and (iv) H9 and H13 were closely related with 2%. The genetic distances between intergroup haplotypes ranged between 8 to 22%.

Phylogenetic clustering of 13 haplotypes showed a similar pattern as seen above. Trees constructed using three different methods showed similar topology (Supplementary **Figure S2**). The haplotypes were majorly grouped into four clusters as identified in genetic distance analysis.

Haplotype network also showed similar patterns of clustering (**Figure 2**). Thirteen haplotypes were portioned into four clusters: Cluster-1 comprising H9 and H13; cluster-2 comprising H8 and H14; cluster-3 comprising H4, H2, H3, H12, H10 and H1; and cluster-4 comprising H6, H11 and H15.

**Figure 2.**
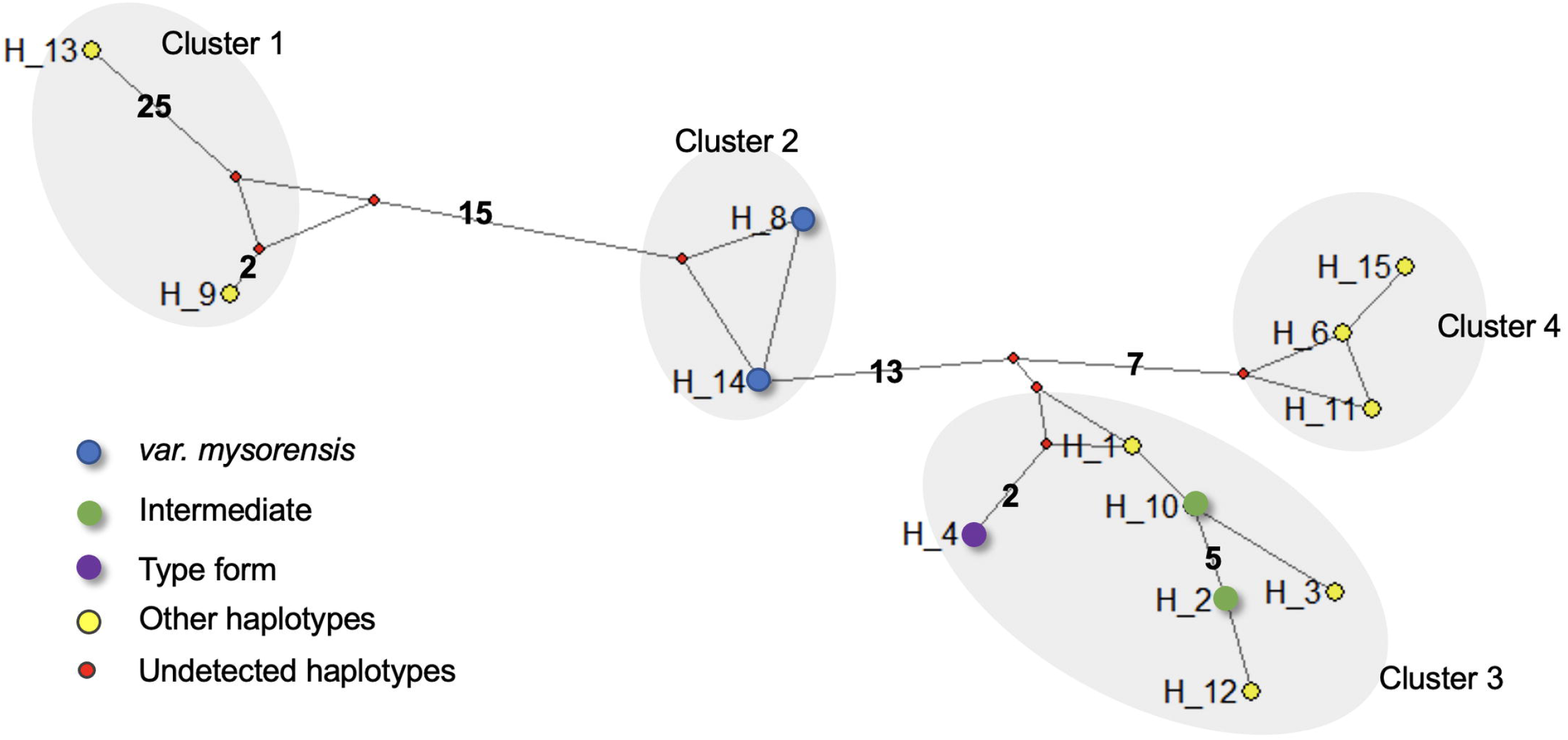
Haplotype network based on the *AsteObp1* intron 1 sequences of *An. stephensi*. Median-oining network was constructed using NETWORK 10.2. Nodes represent haplotypes (assigned as markers for biological form, other haplotypes and undetected haplotypes). Numbers shown on the branches (except one step mutations) represent the numbers of mutations separating each haplotype.

None of the above analyses could partition haplotypes designated as markers for ’ type form’ (H4) and ’ intermediate’ (H2 and H10), which fall within the same cluster. However, the haplotypes designated as markers for *’ var. mysorensis*’ (H8 and H14) from an independent cluster.

### Association of *AsteObp1* haplotype groups with biological forms of *An. stephensi*

According to Firooziyan et al. [23], haplotypes H8 and H14 are assigned as markers for the ‘*var. mysorensis’*, H2 and H10 for the ‘intermediate’ and H4 for the ‘type form’ (Supplementary **Table S1**). Firooziyan et al. [23] considered closely related haplotypes with indel and SNP into a similar category and therefore, haplotypes H1, H3 and H12 being closely related (genetic distances ≤0.2) to haplotype H2 and H10, can also be included in the group of markers for ‘intermediate’. The other haplotypes (H9, H13, H6, H11 and H15), which constitute 25% of the total haplotypes in field populations and 66% in a laboratory colony, could not be classified into any of the above categories, being genetically distantly related to any of the predefined markers (8-22%). To evaluate the association of such markers with biological forms, we, therefore, classified haplotypes into four categories: haplogroup-M (HG-M) for ‘*var. mysorensis’* comprising H8 and H14, (ii) haplogroup-I (HG-I) for ‘intermediate’, comprising H2, H3, H12, H10 and H1, (iii) haplotype-T (H-T) for ‘type form’, i.e., H4, and (iv) other haplotypes (H9, H13, H6, H11 and H15). The frequency distribution of different haplotype groups, categorized above, among phenotypically determined biological forms based on the number of floats on eggs, has been shown in **Figure 3**. Due to overlapping mode numbers of egg ridges defined for biological forms, mosquitoes having mode numbers ≤ 13 were considered ‘*var. mysorensis’*, ≥16 as ‘type form’ and 14-15 as ‘intermediate’. There was no significant difference in the frequencies of different haplotype groups in three biological forms among field populations **Figure 3** based on chi-square test as well as overlapping 95% confidence interval (CI). In another analysis, when we analysed the distribution of haplotype groups among biological forms classified based on mode number of egg ridges ≤ 12, 14-15 and ≥17 (excluding overlapping mode number of intermediate form), similar results were obtained. Interestingly, HG-T was absent from type form and HG-M was absent from ‘*var. mysorensis*’ in the latter analysis. No test was performed between the field population and the laboratory colony. The average pairwise difference between biological forms ranged between 13.58 and 15.21 and within biological forms ranged between 13.59 and 14.19 (**Table S2**). *F*_ST_ values between the three biological forms were negative (−0.007 to -0.014), which is equivalent to zero, and were not significantly different (*p*=0.65), which means there is no genetic subdivision between the populations (**Table S3**) based on *AsObp1* sequences.

**Figure 3.**
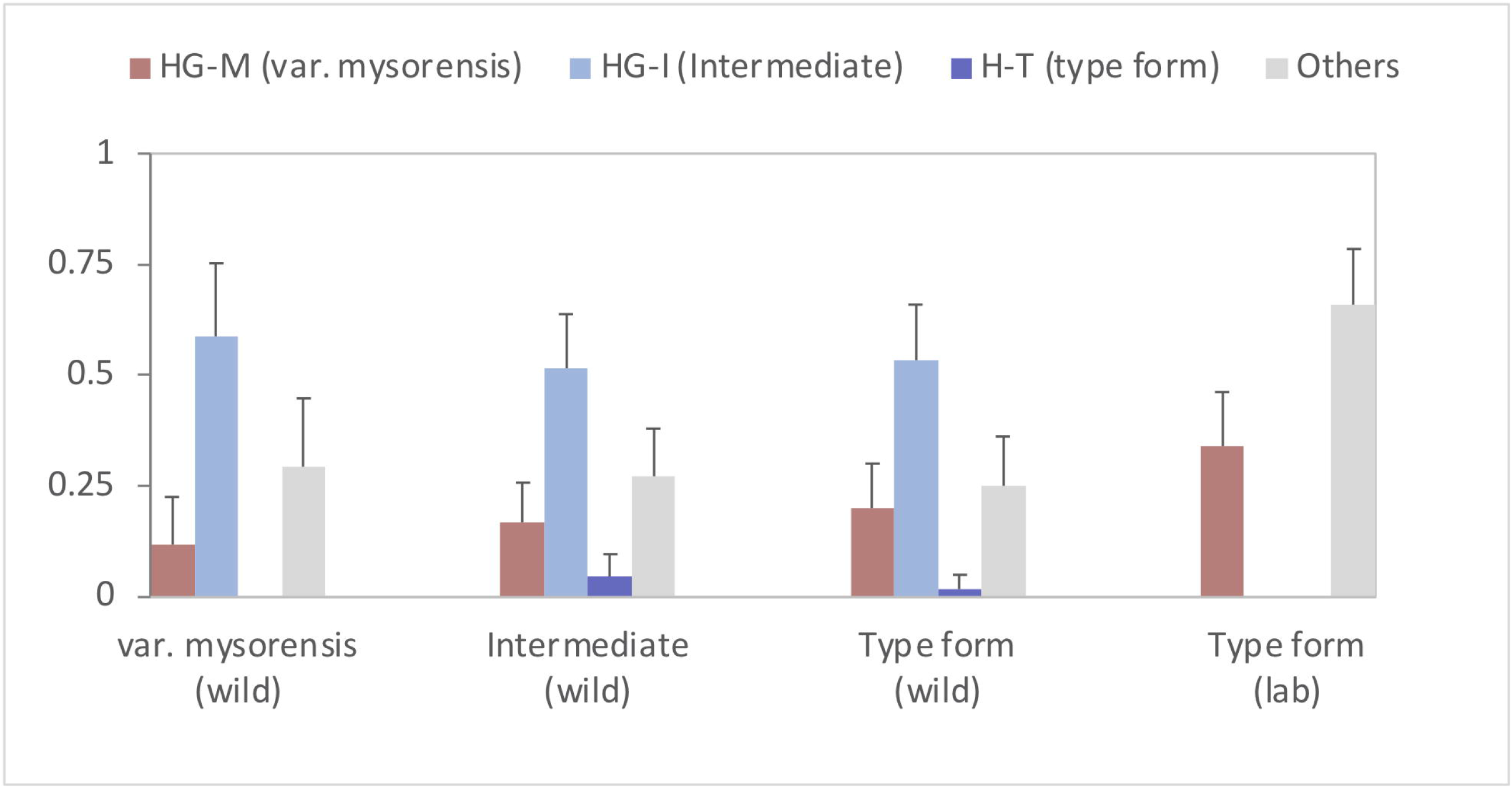
Distribution of haplotype groups in morphologically identified biological forms in field opulations and a laboratory strain. Error bars represent 95% confidential interval (CI).

### Characterization of field and laboratory populations based on egg-ridges and *AstObp1* markers

#### 1. Delhi-colony (Type form)

The laboratory colony of Delhi (NIMR strain) was characterized as a typical ‘type form’ based on floats on egg-ridges. The mode number of ridges ranged between 16 and 20 (average: 18.29). However, the expected haplotype H4 (marker for ‘type form’) was absent. Instead, two different haplotypes, H8 (marker for *var. mysorensis*) and H9 were found with frequencies of 0.34 and 0.66, respectively. These two haplotypes are far distantly related to the designated haplotype for ‘type form’ (H4) with T-3-P distances of 17 and 18%, respectively (**Figure S1)**

#### 2. Bengaluru population (Type form)

The mode number of ridges ranged between 16 and 18 (mean:16.55) which can be categorized as ‘type form’. The expected haplotype H4, the marker for ‘type form’, was not found in this population. The Bengaluru population had haplotypes belonging to HG-I (50%), HG-M (25%) and ‘others’ (25%). Haplogroup H-T was absent.

#### 3. Village Chilawali village (var. mysorensis)

The mode number of ridges in the Chilwali population ranged between 11 and 14 (average: 13.18), which can be categorized as ‘*var. mysorensis*’. Out of a total of 18 haplotypes identified, eight belonged to HG-I, one to HG-T and nine to ‘others’. Expected haplotypes (HG-M) were not found.

#### 4. Villages Karamcand Pur (var. mysorensis and intermediate)

The mode number of ridges in the Karamcand Pur population ranged between 12 and 16 (average: 14.85) which can be categorized as a mixture of ‘intermediate’ and ‘*var. mysorensis*’ forms. The Karamcand Pur population had all the haplotypes identified in this study (**Table 2**) except H12 and H14. 48% of haplotypes belonged to HG-I; 31% to the ’ others’ group; 17% to HG-M and 4% to H-T.

#### 5. Villages Jaisingh Pur (all forms)

The mode number of ridges ranged between 11 and 16 (average 13.69). All forms were present in the village. Majority of haplotypes were belonging to HG-I (66%), followed by HG-M and the ‘others’ group (each 16%) and H-T(2%).

### Association of intron-1 of *AsteObp1* with biological form in different laboratory colonies of *An. stephensi*

The list of haplotypes of intron-1 of *AsteObp1* and their biological forms in different laboratory strains, all originating from the Indian subcontinent, has been listed in **Table 3**. Interestingly, all the six strains had H8 haplotype, a haplotype designated as a marker for ‘*var. mysorensis’*, irrespective of their biological forms determined phenotypically. Delhi strain, which is a relatively new strain (5 years old) had additional haplotype H9, which is not assigned to any biological form. Among all strains, only the Hor strain was characterized as ‘*var. mysorensis*’ morphologically. Two strains; AsteI2 [29] and NIMR strain were identified as ‘type form’ and strain SDA500 [20], as ‘intermediate’ form. The biological form of the remaining two strains (UCISS2018 and STE2) is unknown. Thus, *AstObp1* haplotypes failed to identify biological forms of laboratory colonies.

**Table 3.** *AsObp1-*intron haplotype in different laboratory strains of *An. stephensi* maintained and their biological forms

| Strain (origin) | *AsObp1*-intron haplotype (assigned biological form*) | Reference (sequence accession number) | Biological form | Reference |
| --- | --- | --- | --- | --- |
| 1. AsteI2 (India) | H8 (*var. mysorensis*) | KE389038† | ‘Type’ form | Jiang et al., 2014 [29] |
| 2. SDA500 (Pakistan) | H8 (*var. mysorensis*) | KB664334†, CP032232§ | ‘Intermediate’ | Shinzawa et al., 2013 [20] |
| 3. Hor (India) | H8 (*var. mysorensis*) | MW013512-MW013520§<br>MZ420719-MZ420722§ | ‘*var. mysorensis*’ | Khan et al., 2022 [34] |
| 4. NIMR (Delhi, India) | H8 (*var. mysorensis*)+H9** | This study | ‘Type’ form | This study |
| 5. STE2 (Delhi, India) | H8 (*var. mysorensis*) | This study |  |  |
| 6. UCISS2018 (India) | H8 (*var. mysorensis*) | CM023248† |  |  |

† retrieved from VectorBase; § retrieved from GenBank; \*following Gholizadeh et al. and Firooziyan et al.; \*\*haplotype not assigned to any biological form

## Discussion

Evidence suggests that the biological forms of *An. stephensi* are ecological variants exhibiting differences in their biological characteristics including vectorial competence [19-20]. It is, therefore, vital to identify biological forms in a field population. Previous attempts to identify a molecular marker for the differentiation of ‘type form’ and ‘*var. mysorensis*’ based on ribosomal DNA (ITS2 and 28S rDNA) [30-31] and mitochondrial markers [32-33] were unsuccessful. However, Gholizadeh et al. [22] reported that each of the three biological forms had a fixed *AsteObp1* intron-1 haplotype and proposed it as a molecular marker for their identification. The observation, by Gholizadeh et al. [22], that fixed differences do exist between different biological forms in intron-1 was based on the four inbred laboratory colonies (two of which have completed over 200 generations) representing three biological forms. Very recently, confirmation of the association of an intron haplotype with a biological form (*var. mysorensis*) was shown by the same research group [34] in a highly inbred and old (>38 years) laboratory strain (Hor strain) originating from India. However, the proposed diagnostic markers have never been tested on field mosquitoes that have been characterized for biological forms based on egg morphology. Such exercise was desirable because the fixation of a haplotype in a laboratory colony can be due to the founder effect, which tends to reduce genetic variability and increases homozygosity over generations [35]. A similar effect was seen in the present study, where we found reduced heterozygosity in a laboratory colony (New Delhi) with the presence of just two haplotypes in contrast to field populations that were highly polymorphic. Therefore, any conclusion based on the association of a haplotype with biological form in laboratory colonies is flawed.

In another study by the same group [23], the biological forms were speculated to be sibling species and were designated as sibling species A, B and C. However, in this study, the assignment of biological form was based on the presumed association of haplotype with biological form and the association of haplotypes with biological forms identified based on egg morphology was not established. The limited data suggested the absence of heterozygotes based on sequences from five samples from Afghanistan and 13 samples from Iran. There is no mention of the presence or absence of heterozygotes in the paper. In both the studies by the Gholizadeh group [22-23], large numbers of samples (150 and 100 samples, respectively) were PCR-amplified, but only a few samples (20 and 18, respectively) were sequenced, which may be a reason why heterozygotes were not observed or may have been missed due to ambiguous sequences due to indels.

In the present study, we observed a high degree of heterozygosity in Indian field populations with low inbreeding coefficients indicating unrestricted gene flow. The observed high degree of heterozygosity (>80%) in the field population is not unusual considering the number of haplotypes present in the population. The heterozygosity is expected to increase with the increase in the number of haplotype alleles. With the presence of 13 haplotype alleles, theoretically, we expect maximum heterozygosity up to 0.9 when alleles are equally frequent (H_*E* Max_=1-(1/*n*); see Supplementary Appendix S1 for the derivation). A limited number of sequencing in this study revealed the presence of at least 13 haplotypes of intron-1. The number of haplotypes will keep increasing with the increase in sample size and populations because introns, being neutral, are subjected to a high rate of evolution.

In our study, we did not find an association of any *AstObp1* intron-1 haplotype with a specific biological form. Moreover, we didn’t find haplotype-T (H4) in a laboratory colony morphologically characterized as ‘type form’. Similarly, haplotype group-M (H8 and H14) was absent in a population (Chilwali, Haryana), characterized morphologically as ‘*var. mysorensis*’. Inconsistency in the association of haplotype with biological form was also noted in several laboratory strains, originating from India and Pakistan and being maintained for long (more than 30 years). Interestingly, all such laboratory colonies had H8 haplotype (designated for *mysorensis*) irrespective of their biological forms (**Table 3**).

In this study, we also found low inbreeding coefficients (close to zero) in both, laboratory as well as field populations, based on the *AsteObp1* intron-1, which refutes the presence of putative sibling species based on this marker in Indian populations. Whether reproductive isolation does exist in this species in the Pacific region needs to be confirmed by haplotyping field populations for *AsteObp1* markers and establishing their association with egg morphology. The negative values of *F*_*ST*_ between biological forms (−0.007 to -0.014) based on the *AsObp1* intron-1 sequence indicate lack of genetic subdivision. However, a single marker is not sufficient to draw any conclusion on population structure; but the use of this *AsObp1* marker for the identification of biological forms or putative sibling species may lead to serious consequences.

The present study provides a cautionary note that the *AsteObp1* intron-1 is not a suitable marker for the identification of biological forms or putative species but does not rule out the presence of putative sibling species or genetic constraints in gene flow between the forms. Previous genetic analyses have shown conflicting reports on gene flow between different biological forms occupying different ecological niches. A population genetic analysis on Indian *An. stephensi* populations using microsatellite markers indicated high genetic differentiation (*F*_ST_) and negligible gene flow between the three variants [36]. However, extensive gene flow among the three forms of *An. stephensi* (type form, *var. mysorensis* and intermediate) have been reported in Iran based on mitochondrial genes *cytochrome oxidase 1* and *cytochrome oxidase 2* (*COI* and *COII*) [32-33]. Irrespective of biological forms, the Saudi Arabian *An. stephensi* have been found to be different from all other populations, viz., India, Pakistan, Djibouti, Ethiopia and derived colony strains based on *COI* sequence [37-38]. However, the ITS2 sequence of *An. stephensi* from Saudi Arabia [37] is no different from Indian, Sri Lankan and African populations [31, 39-40], exhibiting mito-nuclear discordance [41]. Though ITS2 has been widely and successfully used for discrimination of species, in some cases, for example in the case of Hyrcanus Complex, failed to differentiate *An. pseudopictus* and *An. hyrcanus* [42]. So far, there is no evidence of differences in ribosomal DNA [30-31]. Therefore, there is a need to extend studies until find good marker for molecular identification of these biological forms.

The objective of this study was to evaluate the utility of *AsObp1* as a marker to identify biological forms of *An. stephensi* but by no means is indented to study the population structure or evolutionary aspects, which remains to be studied using multiple markers. Therefore, there is an urgent need to perform a robust population genetic analysis using multiple molecular markers on morphologically characterized individuals to establish genetic diversity, population structure, and extent of gene flow between different biological forms. Next generation sequencing, on the other hand, may not only provide genetic differentiation among different biological forms; but can also reveal the underlying genetic mechanisms of ecological adaptation of different biological forms.

## Conclusions

*AsteObp1*-intron is not a suitable marker for the identification of biological forms, at least in Indian populations. No specific haplotype was found associated with biological forms. The presence of a high degree of heterozygosity and low inbreeding coefficients in Indian populations refutes the probable existence of sibling species in *An. stephensi* based on *AsteObp1* marker, at least in India.

## Ethics approval and consent to participate

Not applicable

## Consent for publication

Not applicable

## Acknowledgments

Authors are thankful to the Director, NIMR for providing facilities to conduct the work. The technical assistance rendered by Mr. Uday Prakash, Mr. Kanwar Singh and Mr. Sadarudddin is acknowledged. The following reagent was obtained through BEI Resources, NIAID, NIH: Genomic DNA from *Anopheles stephensi*, Strain STE2, MRA-144, contributed by Mark Q. Benedict.

## References

1. Dash AP, Adak T, Raghavendra K, Singh OP. The biology and control of malaria vectors in India. Curr Sci. 2007; 92, 1571–1578.

2. Sharma SK, Hamzakoya KK. Geographical spread of Anopheles stephensi vector of urban Malaria, and Aedes aegypti, vector of dengue/ DHF, in the Arabian Sea Islands of Lakshadweep, India. Dengue Bulletin. 2001; 25: 88–91. https://apps.who.int/iris/handle/10665/148798

3. Gayan Dharmasiri AG, Perera AY, Harishchandra J, Herath H, Aravindan K, Jayasooriya HTR, et al. First record of Anopheles stephensi in Sri Lanka: a potential challenge for prevention of malaria reintroduction. Malar J. 2017; 16:326. doi: 10.1186/s12936-017-1977-7.

4. Faulde MK, Rueda LM, Khaireh BA. First record of the Asian malaria vector Anopheles stephensi and its possible role in the resurgence of malaria in Djibouti, Horn of Africa. Acta Trop. 2014; 139:39–43. doi: 10.1016/j.actatropica.2014.06.016.

5. Carter TE, Yared S, Gebresilassie A, Bonnell V, Damodaran L, Lopez K, et al. First detection of Anopheles stephensi Liston, 1901 (Diptera: Culicidae) in Ethiopia using molecular and morphological approaches. Acta Trop. 2018; 188:180–186. doi: 10.1016/j.actatropica.2018.09.001.

6. WHO. Vector alert: Anopheles stephensi invasion and spread. 26 August 2019. https://www.who.int/news/item/26-08-2019-vector-alert-anopheles-stephensi-invasion-and-spread (retrieved: 1 October, 2021)

7. Seyfarth M, Khaireh BA, Abdi AA, Bouh SM, Faulde MK. Five years following first detection of Anopheles stephensi (Diptera: Culicidae) in Djibouti, Horn of Africa: populations established-malaria emerging. Parasitol Res. 2019; 118:725–732. doi: 10.1007/s00436-019-06213-0.

8. Balkew M, Mumba P, Dengela D, Yohannes G, Getachew D, Yared S, et al. Geographical distribution of Anopheles stephensi in eastern Ethiopia. Parasit Vectors. 2020; 13:35. doi: 10.1186/s13071-020-3904-y.

9. Ahmed A, Khogali R, Elnour MB, Nakao R, Salim B. Emergence of the invasive malaria vector Anopheles stephensi in Khartoum State, Central Sudan. Parasit Vectors. 2021;14(1):511. doi: 10.1186/s13071-021-05026-4.

10. de Santi VP, Khaireh BA, Chiniard T, Pradines B, Taudon N, Larréché S, Mohamed AB, de Laval F, Berger F, Gala F, Mokrane M, Benoit N, Malan L, Abdi AA, Briolant S. Role of Anopheles stephensi Mosquitoes in Malaria Outbreak, Djibouti, 2019. Emerg Infect Dis. 2021; 27:1697–1700. doi: 10.3201/eid2706.204557.

11. Sinka ME, Pironon S, Massey NC, et al. A new malaria vector in Africa: Predicting the expansion range of Anopheles stephensi and identifying the urban populations at risk. Proc Natl Acad Sci U S A. 2020;117(40):24900–24908. doi:10.1073/pnas.2003976117

12. Sweet WC, Rao BA. Races of A. stephensi Liston,1901. Indian Medical Gazette. 1937; 72,665–674.

13. Rao BA, Sweet WC, Subbarao, A.M. Ova measurements of A. stephensi type and A. stephensi var. mysorensis. Journal of the Malaria Institute of India. 1938; 1,261–266.

14. Subbarao SK, Vasantha K, Adak T, Sharma VP, Curtis CF. Egg-float ridge number in Anopheles stephensi: ecological variation and genetic analysis. Med Vet Entomol. 1987; 1: 265–71. doi: 10.1111/j.1365-2915.1987.tb00353.x.

15. Sweet W, Rao B, Rao A. Cross-breeding of An. stephensi type and An. stephensi var. mysorensis. J Malar Inst India. 1938;1:149–54.

16. Rutledge LC, Ward RA, Bickley WE. Experimental hybridization of geographic strains of Anopheles stephensi (Diptera: Culicidae). Ann Entomol Soc Am. 1970; 63:1024–30.

17. Manouchehri AV, Javadian E, Eshighy N, Motabar M. Ecology of Anopheles stephensi Liston in southern Iran. Trop Geogr Med. 1976;28(3):228–232.

18. Oshaghi MA, Yaaghoobi F, Vatandoost H, Abaei MR. Anopheles stephensi biological forms: geographical distribution and malaria transmission in malarious region of Iran. Pak J Biol Sci. 2006a; 9: 294–6.

19. Basseri HR, Mohamadzadeh Hajipirloo H, Mohammadi Bavani M, Whitten MM. Comparative susceptibility of different biological forms of Anopheles stephensi to Plasmodium berghei ANKA strain. PLoS One. 2013; 8:e75413. doi: 10.1371/journal.pone.0075413.

20. Shinzawa N, Ishino T, Tachibana M, Tsuboi T, Torii M. Phenotypic dissection of a Plasmodium-refractory strain of malaria vector Anopheles stephensi: the reduced susceptibility to P. berghei and P. yoelii. PLoS One. 2013; 8: e63753. doi: 10.1371/journal.pone.0063753.

21. Nagpal BN, Srivastava A, Kalra NL, Subbarao SK. Spiracular indices in Anopheles stephensi: a taxonomic tool to identify ecological variants. J Med Entomol. 2003;40: 747–9. doi: 10.1603/0022-2585-40.6.747.

22. Gholizadeh S, Firooziyan S, Ladonni H, Hajipirloo HM, Djadid ND, Hosseini A, Raz A. The Anopheles stephensi odorant binding protein 1 (AsteObp1) gene: a new molecular marker for biological forms diagnosis. Acta Trop. 2015; 146:101–13. doi: 10.1016/j.actatropica.2015.03.012.

23. Firooziyan S, Dinparast Djadid N, Gholizadeh S. Speculation on the possibility for introducing Anopheles stephensi as a species complex: preliminary evidence based on odorant binding protein 1 intron I sequence. Malar J. 2018; 17:366. doi: 10.1186/s12936-018-2523-y

24. Christophers SR. The fauna of British India, including Ceylon and Burma. Diptera. 4. Family Culicidae, Tribe Anophelini. London: Tribe Anophelini. Taylor and Francis; 1933

25. Black WC and Duteau NM. The molecular biology of insect disease vectors: a methods manual. In: RAPD-PCR and SSCP analysis for insect population genetic studies (Crampton, J.M., Beard, C.B. & Louis, C., eds.). Chapman & Hall, London, 1997; 361–373.

26. Hill JT, Demarest BL, Bisgrove BW, Su YC, Smith M, Yost HJ. Poly peak parser: Method and software for identification of unknown indels using sanger sequencing of polymerase chain reaction products. Dev Dyn. 2014;243:1632–6. doi: 10.1002/dvdy.24183.

27. Kumar S, Stecher G, Tamura K. MEGA7: Molecular Evolutionary Genetics Analysis Version 7.0 for Bigger Datasets. Mol Biol Evol. 2016; 33:1870–4. doi: 10.1093/molbev/msw054.

28. Excoffier L, Lischer HE. Arlequin suite ver 3.5: a new series of programs to perform population genetics analyses under Linux and Windows. Mol Ecol Resour. 2010;10(3):564–7. doi: 10.1111/j.1755-0998.2010.02847.x

29. Jiang X, Peery A, Hall AB, Sharma A, Chen XG, Waterhouse RM, Komissarov A, Riehle MM, Shouche Y, Sharakhova MV, Lawson D, Pakpour N, Arensburger P, Davidson VL, Eiglmeier K, Emrich S, George P, Kennedy RC, Mane SP, Maslen G, Oringanje C, Qi Y, Settlage R, Tojo M, Tubio JM, Unger MF, Wang B, Vernick KD, Ribeiro JM, James AA, Michel K, Riehle MA, Luckhart S, Sharakhov IV, Tu Z. Genome analysis of a major urban malaria vector mosquito, Anopheles stephensi. Genome Biol. 2014;15(9):459. doi: 10.1186/s13059-014-0459-2.

30. Alam MT, Bora H, Das MK, Sharma YD. The type and mysorensis forms of the Anopheles stephensi (Diptera: Culicidae) in India exhibit identical ribosomal DNA ITS2 and domain-3 sequences. Parasitol Res. 2008;103:75–80. doi: 10.1007/s00436-008-0930-7.

31. Mishra S, Sharma G, Das MK, Pande V, Singh OP. Intragenomic sequence variations in the second internal transcribed spacer (ITS2) ribosomal DNA of the malaria vector Anopheles stephensi. PLoS One. 2021; 16: e0253173. doi: 10.1371/journal.pone.0253173.

32. Chavshin AR, Oshaghi MA, Vatandoost H, Hanafi-Bojd AA, Raeisi A, Nikpoor F. Molecular characterization, biological forms and sporozoite rate of Anopheles stephensi in southern Iran. Asian Pac J Trop Biomed. 2014; 4: 47–51. doi:10.1016/S2221-1691(14)60207-0

33. Oshaghi MA, Yaaghoobi F, Abaie M. Pattern of mitochondrial DNA variation between and within Anopheles stephensi (Diptera: Culicidae) biological forms suggests extensive gene flow. Acta Trop. 2006b; 99: 226–233

34. Khan J, Gholizadeh S, Zhang D, Wang G, Guo Y, Zheng X, Wu Z, Wu Y. Identification of a biological form in the Anopheles stephensi laboratory colony using the odorant-binding protein 1 intron I sequence. PLoS One. 2022 Feb 22;17(2):e0263836. doi: 10.1371/journal.pone.0263836.

35. Moorad JA, Wade MJ. A genetic interpretation of the variation in inbreeding depression. Genetics. 2005;170: 1373–84. doi: 10.1534/genetics.104.033373.

36. Vipin, Dube M, Gakhar SK. Genetic differentiation between three ecological variants (‘type’, ‘mysorensis’ and ‘intermediate’) of malaria vector Anopheles stephensi (Diptera: Culicidae) Insect Science. 2010; 17: 335–343, doi 10.1111/j.1744-7917.2010.01316.x

37. Munawar K, Saleh A, Afzal M, et al. Molecular characterization and phylogenetic analysis of anopheline (Anophelinae: Culicidae) mosquitoes of the Oriental and Afrotropical Zoogeographic zones in Saudi Arabia. Acta Trop. 2020;207:105494. doi:10.1016/j.actatropica.2020.105494

38. WRBU. Anopheles stephensi Liston, 1901. https://www.wrbu.si.edu/vectorspecies/mosquitoes/stephensi (retrieved 8 November, 2021)

39. Surendran SN, Sivabalakrishnan K, Gajapathy K, Arthiyan S, Jayadas TTP, Karvannan K, et al. Genotype and biotype of invasive Anopheles stephensi in Mannar Island of Sri Lanka. Parasit Vectors. 2018; 11:3. doi: 10.1186/s13071-017-2601-y.

40. Tadesse FG, Ashine T, Teka H, et al. Anopheles stephensi mosquitoes as vectors of Plasmodium vivax and falciparum, Horn of Africa, 2019. Emerg Infect Dis. 2021;27(2):603–607. doi:10.3201/eid2702.200019

41. Singh OP, Sindhania A, Sharma G, et al. Are members of the Anopheles fluviatilis complex conspecific?. Acta Trop. 2021;224:106149. doi:10.1016/j.actatropica.2021.106149

42. Fang Y, Shi WQ, Zhang Y. Molecular phylogeny of Anopheles hyrcanus group members based on ITS2 rDNA. Parasit Vectors. 2017;10(1):417. doi:10.1186/s13071-017-2351-x

