## Supplementary material for "Evaluation of intron-1 of odorant-binding protein-1 of *Anopheles stephensi* as a marker for the identification of biological forms or putative sibling species": Table S1

**Table S1** List of GenBank submission designated for the identification of biological forms in *An. stephensi* by Gholizadeh et al. and Firooziyan et al., and designation of identical haplotype in this study

| Marker defined for the biological form | GenBank accession numbers | References | Haplotype designation in this study |
|---|---|---|---|
| *var. mysorensis* | KJ557449—KJ557451, KJ557453—KJ557455, KJ557457—KJ557461, KJ557466—KJ557467 | Gholizadeh et al.; 2015 | H8 |
| *var. mysorensis* | KT587049, KT587051 | Firooziyan et al.; 2018 | H14 |
| Intermediate | MG797534 | Firooziyan et al.; 2018 | H2 |
| Intermediate | KJ557452, KJ557456, KJ557464, KJ557468 | Gholizadeh et al.; 2015 | H2 |
| Intermediate | KT587050, KT587052- KT587053 | Firooziyan et al.; 2018 | H10 |
| Type form | KJ557462—KJ557463, KJ557465 | Gholizadeh et al.; 2015 | H4 |
| Type form | MG797525—MG797533 | Firooziyan et al.; 2018 | H4 |
