## Supplementary material for "Evaluation of intron-1 of odorant-binding protein-1 of *Anopheles stephensi* as a marker for the identification of biological forms or putative sibling species": Table S2

**Table S2** Pairwise $F_{ST}$ values between the three biological forms based on *AsObp1*-intron-1 sequences

| | Type form | Intermediate | Mysorensis |
|---|---|---|---|
| Type form | 0 | | |
| Intermediate | -0.00691 | 0 | |
| Mysorensis | -0.01388 | -0.01392 | 0 |

*p* values non-significant
