## Supplementary material for "Evaluation of intron-1 of odorant-binding protein-1 of *Anopheles stephensi* as a marker for the identification of biological forms or putative sibling species": Table S3

**Table S3** Average pairwise differences between and within biological forms. Values above diagonal are pairwise differences between populations (PiXY), diagonal elements are pairwise differences within population (PiX) and below diagonal are corrected pairwise difference (PiXY-(PiX+PiY)/2)

| | Type form | Intermediate | Mysorensis |
|---|---|---|---|
| Type form | 13.58983 | 13.77576 | 14.19657 |
| Intermediate | -0.09492 | 14.15152 | 14.47504 |
| Mysorensis | -0.20441 | -0.20677 | 15.21212 |

*p* values non-significant
