## Supplementary Appendix 1 for "Evaluation of intron-1 of odorant-binding protein-1 of *Anopheles stephensi* as a marker for the identification of biological forms or putative sibling species"

**Supplementary appendix: S1**

Expected heterozygosity ($H_E$) for n alleles

$H_E = 1-\sum_{i=1}^{n}(pi)\sum_{i=1}^{n}(pi)^2$

(Where *pi* = frequency of ith allele of n alleles)

Maximum expected heterozygosity ($H_{E\_Max}$) will be when alleles are equally frequent (i.e., *pi* = 1/n) in a population

Then,

$H_{E\_Max} = 1-(n(1/n)^2)$

$= 1-(1/n)$
