## Supplementary material for "Evaluation of intron-1 of odorant-binding protein-1 of *Anopheles stephensi* as a marker for the identification of biological forms or putative sibling species": Figure S1

| | H1 | H10 | H2 | H3 | H12 | H4 | H6 | H11 | H15 | H8 | H14 | H9 | H13 |
|---|---|---|---|---|---|---|---|---|---|---|---|---|---|
| H1 | | 1 | 1 | 2 | 2 | 3 | 9 | 9 | 10 | 16 | 15 | 16 | 14 |
| H10 | 0.01 | | 0 | 1 | 1 | 4 | 10 | 10 | 11 | 17 | 16 | 17 | 14 |
| H2 | 0.01 | 0 | | 1 | 1 | 4 | 9 | 9 | 10 | 17 | 16 | 15 | 12 |
| H3 | 0.02 | 0.01 | 0.01 | | 2 | 5 | 11 | 11 | 12 | 18 | 17 | 18 | 14 |
| H12 | 0.02 | 0.01 | 0.01 | 0.02 | | 5 | 10 | 10 | 11 | 16 | 15 | 14 | 11 |
| H4 | 0.03 | 0.03 | 0.04 | 0.04 | 0.04 | | 11 | 11 | 12 | 18 | 17 | 18 | 16 |
| H6 | 0.08 | 0.09 | 0.08 | 0.10 | 0.09 | 0.10 | | 1 | 1 | 19 | 18 | 19 | 16 |
| H11 | 0.08 | 0.09 | 0.08 | 0.10 | 0.09 | 0.10 | 0.01 | | 2 | 20 | 19 | 19 | 16 |
| H15 | 0.09 | 0.10 | 0.09 | 0.11 | 0.10 | 0.11 | 0.01 | 0.02 | | 20 | 19 | 20 | 17 |
| H8 | 0.15 | 0.16 | 0.16 | 0.17 | 0.15 | 0.17 | 0.18 | 0.19 | 0.19 | | 1 | 13 | 9 |
| H14 | 0.14 | 0.15 | 0.15 | 0.16 | 0.14 | 0.16 | 0.17 | 0.18 | 0.18 | 0.01 | | 12 | 8 |
| H9 | 0.16 | 0.17 | 0.15 | 0.18 | 0.14 | 0.18 | 0.19 | 0.19 | 0.20 | 0.12 | 0.11 | | 2 |
| H13 | 0.17 | 0.17 | 0.16 | 0.17 | 0.14 | 0.20 | 0.21 | 0.21 | 0.22 | 0.11 | 0.09 | 0.02 | |

**Figure S1** Estimates of Evolutionary Divergence between *AsteObp1* intron-1 haplotypes using the T3P model. The numbers shown above diagonal represent base differences and below diagonal represent genetic distances.
