## Supplementary material for "Evaluation of intron-1 of odorant-binding protein-1 of *Anopheles stephensi* as a marker for the identification of biological forms or putative sibling species": Figure S2

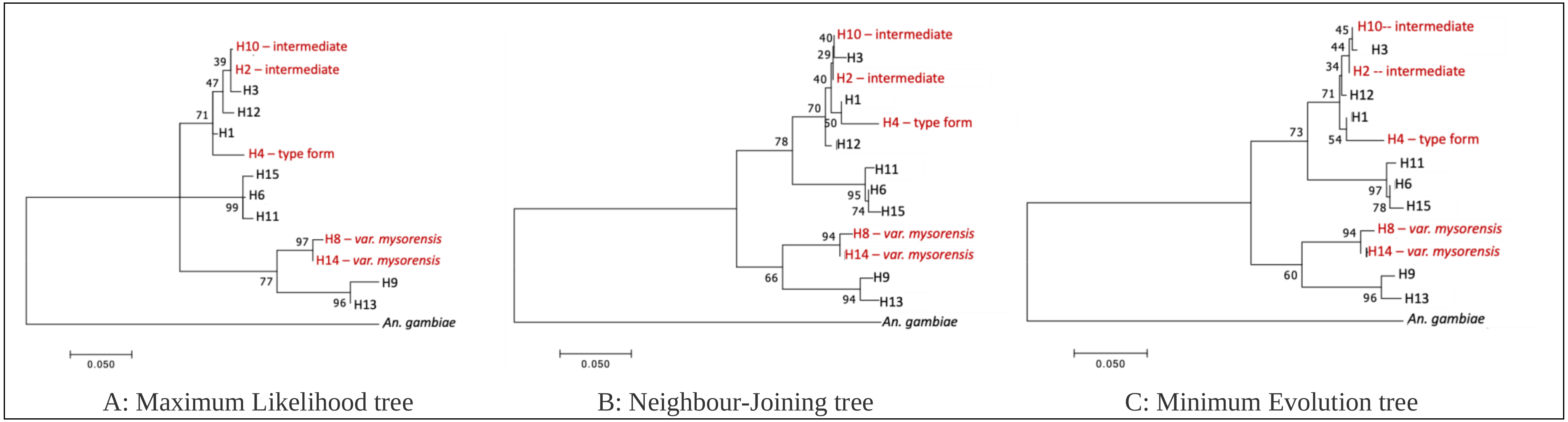


**Figure S2** Phylogenetic tree analysis of *AstObp1* intron haplotypes by Maximum Likelihood, Neighbour-Joining and Minimum-Evolution methods based on T92 model (Tamura 3-parameter), the best model as determined based on lowest Bayesian Information Criterion (BIC) scores. The value on each node represent bootstrap value. The haplotypes labelled with red colour font are markers assigned for the identification of haplotypes by earlier workers. The corresponding intron of *An. gambiae* was taken as outgroup.
